# SEXUAL-DIMORPHISM IN VISUALLY GUIDED APPROACH BEHAVIOR EMERGES UNIQUELY DURING ADOLESCENCE

**DOI:** 10.1101/2024.10.05.616154

**Authors:** Rocio Gonzalez-Olvera, Kelsey Allen, Ting Feng, Jennifer L. Hoy

## Abstract

Studying prey capture behavior in mice offers a fruitful platform for understanding how ecologically relevant visual stimuli are differentially processed by the brain throughout life. For example, specific visual stimuli that indicate prey and that naturally draw appetitive orienting in the adult may be interpreted differently or evoke distinct behaviors during development. What are the neural mechanisms that might allow an animal to flexibly couple the same stimulus information to distinct behavioral outcomes as a function of developmental needs? To begin to address this important question, we quantified visually evoked orienting behaviors between adolescent and adult C57BL6/J mice of both sexes under the natural prey capture context compared to responses in our established virtual motion stimulus paradigm, C-SPOT. Most surprisingly, we discovered that female versus male adolescent mice have opposing innate behavioral biases evoked by the same sweeping motion stimuli which is not present in adulthood. Further, female mice display a robust enhancement of approach towards visual motion over all other groups tested, yet they are the least aggressive in response to live prey. Thus, our work overall revealed that innate approach towards visual motion is robustly dissociated from predatory aggression in female versus male mice specifically during adolescence. This underscores the different natural behavioral goals and physiological states that uniquely apply to female versus male adolescent animals, and reveals that approach towards visual motion is a key sensory-motor process selectively augmented during female adolescence.

**Highlights:**

- Adolescent mice of both sexes innately approach insects more than adults
- Adolescent males versus females have distinct response biases to visual motion
- Adolescent males display the strongest hunger-related predatory aggression

## Introduction

Prey capture is a ubiquitous behavior where animals integrate specific sensory cues, motivational states and motor actions to successfully procure highly rewarding food^1^. Under natural conditions, most animals successfully utilize a range of specific multisensory cues to reliably recognize and capture natural prey^1,2^. Specific prey “cues” are therefore considered a key part of an organism’s sensory ecology that have shaped the evolution of the structure and function of their senses over time^3^. Prey capture behaviors and underlying mechanisms are thus highly conserved across species ^4,5^.

Many species rely on vision in particular to detect, identify and localize palatable prey^6^. There is behavioral, cellular and physiological evidence that sensitivity to prey motion specifically has driven the evolution of eyes and visuo-motor behavior throughout the mammalian lineage^3,6^. Recent studies in the mouse have revealed specialized circuit pathways that link specific motion cues to positive orienting behaviors that ultimately facilitate effective prey foraging behavior in adults^7^, validating a high conservation of functional organization of visual pathways across species in this natural context^7–10^.

The strong conservation of visually-guided prey capture behaviors and use of similar visual cues across species implies a relative “hard-wiring” of underlying neural circuits linking stimulus to response^7–10^. Yet, there is ample evidence of context-dependent regulation of prey-related visual orienting responses and function of underlying neural circuits throughout the animal kingdom. In zebrafish, hunger state robustly alters innate behavioral and underlying neuronal responses to moving visual objects^11^. Insect-naïve adult mice may either avoid or attack live insects without food-deprivation^12,13^, and show more consistent predatory aggression once food-deprivation begins^4^. Indeed, successful prey capture experience reinforces approach responses towards both prey and visual object motion, reducing variability in behavioral responses^4,14^. Similarly, “small object” motion stimuli presented to a nocturnal species of primate facilitates accurate spatial orienting responses in both appetitive and aversive contexts^15^. Thus, multiple agents of selection appear to have reinforced the conservation of prioritizing and responding to the presence of small moving objects in the visual field, while allowing for context-dependent flexibility in an animal’s overt behavioral response to that motion.

Studying the ontogeny of responses to visual motion as it relates to prey pursuit behavior in the mouse presents an opportunity to identify novel developmentally controlled mechanisms that flexibly gate visually driven orienting responses in the mammal^14,16^. Mammals go through several life stages with distinct physiological needs that exert control over dietary requirements^17^. Thus, the immediate value of pursuing prey/small moving objects during development may be distinct from that present in the adult. As adults, mice and other mammals flexibly utilize specific sensory cues to optimize food foraging strategies^18^. However, independent and successful foraging behavior must first emerge robustly after fully weaning off of mother’s milk and near the onset of adolescence in mice, starting at postnatal day 30 (P30). While the onset of adolescence is associated with a wide variety of change in adaptive behaviors, intense competition for food between conspecifics is one of the most critical environmental forces facing newly adolescent weanlings^19^. For example, the territorial dispersal that requires advanced navigation skills to facilitate the establishment of new breeding territory and breeding behaviors themselves do not emerge until late or post adolescence in mice^20^. Moreover, it has been documented in lab strains of house mice, that adolescents consume more food each day than young adult mice^21^. We thus reasoned that mice could be highly tuned towards visual cues indicating nutritious insect prey in early adolescence. In addition, late juvenile, early adolescence is when mammals such as domestic cats engage most readily in approach and attack behavior triggered by motion stimuli that may subserve predation in adulthood^22^. These robust developmental changes characteristic of adolescence could enable a suite of sensory processing and behavioral shifts that facilitate effective and independent prey foraging to offset the potential costs related to increasing their risk of predation.

To test this idea, we quantified cricket capture and visual orienting evoked by computer generated visual motion stimuli in adult and adolescent mice of both sexes^14^. We indeed found robust interactions between sex and age in how mice respond to live cricket prey and visual sweeping motion cues. Adolescent mice of both sexes approached and stayed in proximity to live insect prey more often than adults without attacking as often. They also suppressed or rebounded more quickly from rapid escape-like behavior after initial contact with prey relative to cricket-naïve adults^13^. Most surprisingly, female and male adolescent mice displayed opposing biases in orienting towards visual motion both upon initial detection and subsequent interactions uniquely during this stage of development. While female adolescent mice approached prey and visual motion stimuli more than all other groups, they were the least likely to exhibit predatory aggression towards live crickets. On the other hand, the adolescent males were more likely to persist in arresting in response to visual motion and exhibit more predatory aggression once food restriction began than all other groups. Our work therefore revealed unexpectedly for the first time that a natural sexual dimorphism in visual orienting arises during adolescence in mice. Further, we provide evidence that predatory aggression and approaches towards visual motion are dissociated during adolescence, similar to other mammalian species^22^. This dissociation may provide clues as to the possible underlying hormonal states that give rise to these behavioral correlations^23,24^. Thus, sexually dimorphic behavioral goals and physiological states in adolescent mice lead to significant differences in “innate” orienting responses towards natural prey species and visual object motion.

## Results

### Sex and age interact to yield distinct differences in visual orienting versus predatory aggression towards crickets

Given key changes in adolescent physiology that may serve to motivate food foraging behaviors, we hypothesized that responses towards “prey-like” stimuli would be heightened in adolescent mice (**Fig 1A**). To test this, we designed a 2 × 2 factorial study to determine the relationship between age (factor 1, two ages: P30-45 versus P60-78) and sex (factor 2, two sexes: female versus male) on measured prey capture-related behaviors (**Fig 1**). We found several significant main effects of age and sex and some interactions between sex and age. First, adolescent female mice as a group approach crickets sooner in the first trial (**Fig 1B**) and more frequently through the duration of the first 5-minute trial (**Fig 1C**) than all other groups. Adolescent males also approach before mature adults of both sexes and exhibit more approaches overall than the adults (**see Supplemental Videos 1-4**). Adolescent mice also spent significantly more time near the cricket over the course of the whole 5-minute trial relative to the adults and no main effect of sex was found (**Fig 1D**). The relative position of the cricket in the visual field just prior to approach (**Fig 1E & F**) was not significantly different by age or sex, and, once approaches started, all groups were similarly accurate in their ability to intercept crickets (**Fig 1G**). Taken together, a mouse’s sex, age and the interaction between the two, differentially impacted distinct aspects of prey capture-related behaviors. The strongest effect was seen in an interaction between sex and age for female adolescent mice which responded most quickly to insects and most robustly displayed approaches towards crickets. More direct measures of visual processing related to specific stimulus features, such as relative prey size and position in the visual field were not robustly different by age nor sex (Figure 1 and data not shown).

**Fig. 1.**
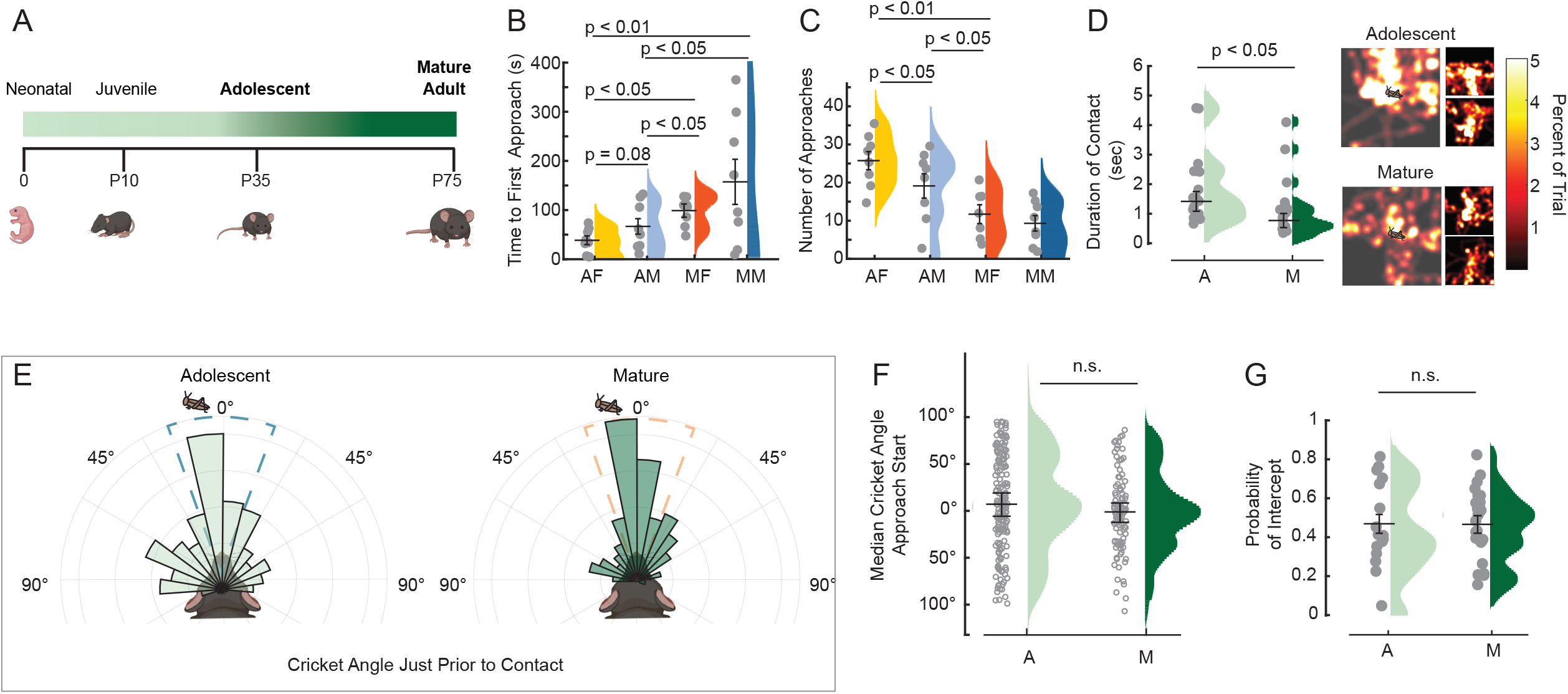
Measures of visual orienting behavior during live prey capture in adolescent mice (P30-45), versus mature mice (>P60) of both sexes during first encounter with crickets. (**A**) Major life-stage classification of mice from postnatal day 0 (P0) through adulthood (P75). The beginning of adolescence for mice is near P30, peaking near P35, the age range of the adolescents in our study. Darkening green indicates maturation and corresponds with data grouped by developmental stage in D-G. (**B**) Average time to first approach for each individual animal (gray circles) on the first day of the first exposure to a cricket for 5 minutes (naive and sated mice). Group distributions are color coded by developmental stage (light shading = adolescents, darker sharding = adults) and sex (warm colors = females, cool colors = males). A = adolescent, M = Mature, F= female, M = male. (**C**) The average number of approaches per animal and group. (**D**) Duration of contact data are plotted by individual, gray circles, and by age group, light versus dark green, adolescent versus mature adults, respectively. No effect of sex, nor a sex by age interaction was found in a two-way ANOVA analysis of duration of contact with cricket during first exposure. Heat maps of average relative location of crickets during exposure in the first 5 minute trial are shown to the right. The large maps are group averages, while the smaller images to the right are representative individuals from each group, female at top and male at bottom (see **Supplemental Videos 1-4**). N=8, 8, 7 & 8 mice, AF, AM, MF and MM groups, respectively; Significance tested for using two-way ANOVA, sex by age, with Tukey’s HSD post hoc testing for identifying significant differences and to correct for multiple comparisons. Normality of each dataset was assessed using the Shapiro-Wilk test, Means shown with overall distribution, Error bars are +/-standard error of the mean.

To further understand how sex and age are related to hunting perse vs. more general exploratory behaviors, we also measured and compared direct measures of predatory aggression including percent of trials with bite attacks of crickets and how long it took mice to capture and kill crickets across 6 days with exposure to 4 crickets per day (**Fig 2**). There was a main effect of age on the percent of trials with attacks on the first day of cricket exposure with both adolescent groups (male and female) displaying fewer attacks relative to the adults (**Fig 2A**). On day two, we discovered a significant interaction between age and sex, with male adolescent mice displaying significantly more attacks than all other groups, while the adolescent females were similar in attack likelihood as both adult sexes (**Fig 2A**). Relatedly, these attack characteristics translated to no captures for adolescent female mice on the first day out of 4 separate cricket trials, while the other groups started to capture insects on the first day, with the adult hunters of both sexes outperforming the adolescent males. However, by day two, the most efficient hunting group were the adolescent males, which quickly turned orienting and approach into attack and kill. The adolescent females, while showing the most rapid responses to crickets and most approaches over a 5-minute period, were slower to attack and kill across the first three days of cricket exposure (**Fig 2B**). Adolescent male mice responded most strongly to the effects of food deprivation, attacking nearly all offered crickets on the first day of food restriction (Day 2). The other groups displayed a more gradual response to food deprivation in measures of lethal attacking (**Fig 2B**). There were significant, but weaker sex-driven differences in adolescent mice in likelihood to attack on the first day of exposure. This baseline performance is measured before food deprivation started (**Fig 2A**, Day 1). After overnight food restriction began, the adolescent male mice clearly lead all groups in exhibiting lethal attacks on all offered insects and reaching peak capture time performance. Our results therefore demonstrate that age by sex interactions determine whether mice turn approach responses into attack and consumptive behaviors. Interest in insects is dissociable from predation perse, in that adolescent females showed the strongest dissociation between detection, approach and predatory attacks. Altogether, this suggests that during adolescence, male and female mice show the most sexual dimorphism in motivational processes related to engaging insects. Males displayed behaviors that indicated more predatory aggression, especially once food deprivation began, while females displayed the most robust approach orienting, yet with reduced aggressive behavior not as readily modified by food restriction until days 4-6.

**Fig. 2.**
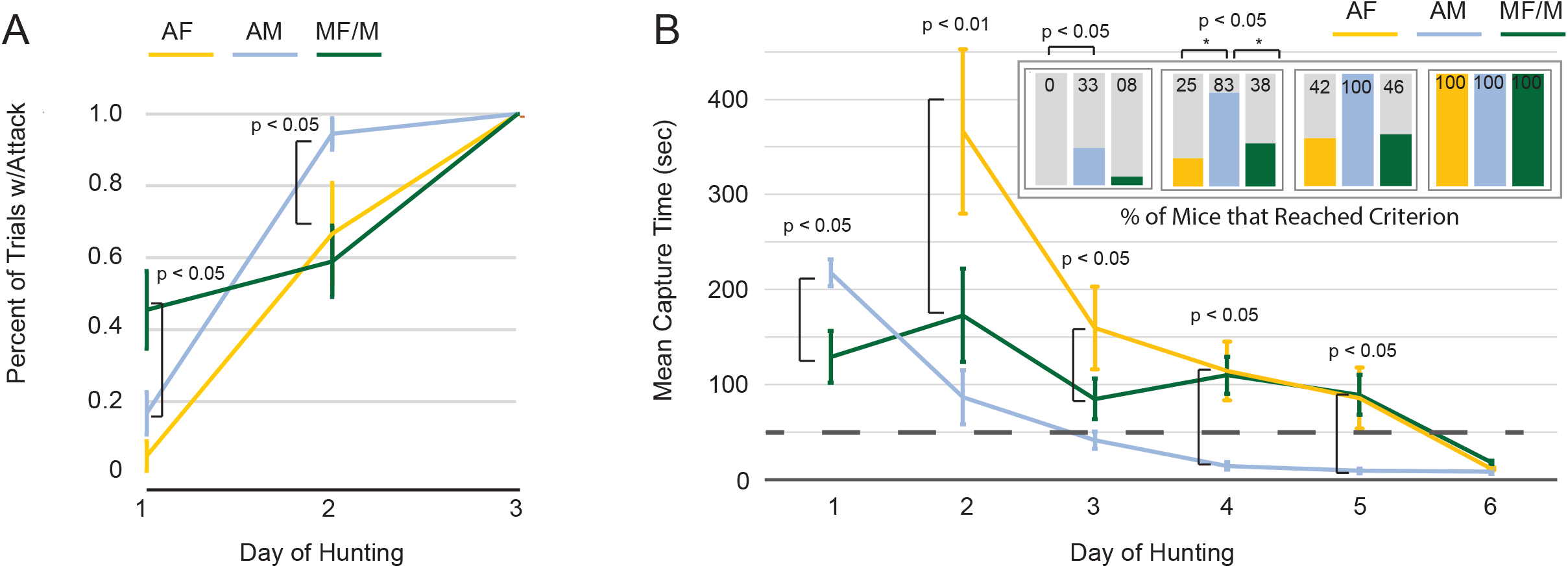
Predatory attack behavior and capture success across 6 days of cricket hunting trials motivated by feeding restriction. (**A**) Percent of trials per day w/attack response. Each mouse was exposed to 1 cricket at a time for up to 5 minutes constituting one trial. Each mouse then experienced up to 4, 5-minute trials per day. (**B**) Mean capture time. If a mouse captured and killed a cricket on a given trial on a given day, their capture time was averaged. The adolescent female mice did not capture any crickets on the first day. Top right insets represent the percentage of mice on each training day (day 2-6) that reached criterion, which is to capture crickets on average in less than 60 seconds. No animals reached criterion on the first day after food restriction began (day 2 on the plot). Significance tested for using a two-way, repeated measures ANOVA, sex x age and the repeated measure over days of training, with Tukey’s HSD post hoc testing for identifying significant differences and to correct for multiple comparisons. N=8, 8, 7 & 8 mice, AF, AM, MF and MM groups, respectively. Adult groups were combined as no effect of sex in adults were found. Normality of each dataset was assessed using the Shapiro-Wilk test, Means shown with overall distribution, Error bars are +/-standard error of the mean.

### Sex and age interact to yield distinct differences in innate visual orienting towards artificial sweeping motion cues

Predation is a complex behavior involving multiple sensory modalities, motivations and cross-modal sensory integration. We therefore sought to understand whether adolescent male and female mice displayed similar orienting responses to visual motion cues that can be utilized during prey capture. We employed our computerized, spontaneous perception of objects task, C-SPOT to detect and quantify innate orienting bias as a function of age and sex in mice^16^. As done previously, we presented a sweeping stimulus along the bottom of the screen positioned as one of the walls of the arena for a total of 60 seconds to each mouse. We focused our analysis on orienting responses to motion stimuli shown previously to evoke the most approaches and fewest arrests in adult mice^14^. Overall, we found main effects of sex and age as well as specific sex and age interactions in how mice responded to the sweeping visual motion stimuli (**Fig 3**). Consistent with observations made when mice reacted to live prey, female adolescent mice, robustly approached sweeping motion stimuli moving at speeds comparable to the movement of live insects (between 2-15 cm/sec real world object speed, **Fig 3A, B**, green box highlighted, **D & E**) more than all the other groups tested. In contrast, the male adolescent mice were more likely to exhibit arrest responses in reaction to the same motion stimuli relative to all the other groups (**Fig 3A, C**, red box highlighted). Despite the sexual dimorphism in how they respond to the motion stimuli, adolescent mice of both sexes responded more and faster overall to the lower field seeping visual motion stimuli than both sexes of adults (**Fig 3D & E, see Supplemental Videos 5-8**). We thus discovered that mice exhibit an innate bias in how they respond to sweeping visual motion objects of a specific size that is specially amplified in adolescence. This is an unexpected and novel observation that indicates the two sexes of mice go through a distinct physiological state during adolescence before they reach a more homogenous adult-like behavioral state in how they respond to both live prey and visual object motion.

**Fig. 3.**
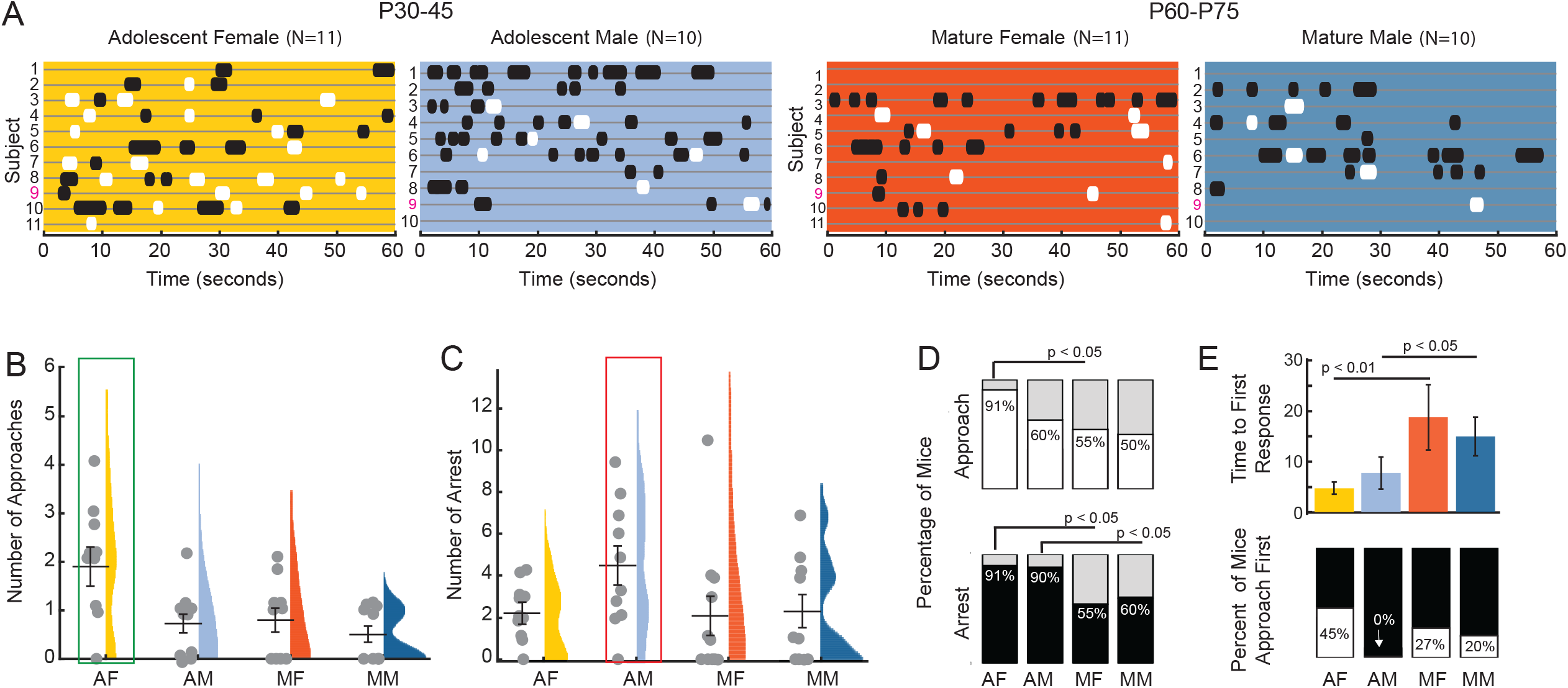
Innate orienting responses to sweeping visual motion in the lower field. (**A**) Ethograms showing the response of all four groups of mice to sweeping visual motion stimuli in the lower environment. Each row (grey line) of the ethogram is the response of an individual animal, white indicates time when animals are approaching stimulus, black is when they arrest in response to the stimulus. The magenta “9” highlights the ethogram corresponding to **supplemental videos 5-8**. (**B**) Mean number of approaches. Green box denotes significant difference in approach number of female adolescents. (**C**) Mean number of arrests, red box denotes significant increase in arrest response in adolescent male mice. (**D**) Percentage of mice in each group exhibiting at least one approach towards sweeping stimulus, white, and those that exhibit at least one arrest in response to the stimulus, black. (**E**) Mean time to first response indicating their sensitivity to the stimulus at top, and likelihood that they approach the stimulus before arresting. Significance tested for using a two-way, ANOVA, sex by age, with Tukey’s HSD post hoc testing for identifying significant differences and to correct for multiple comparisons. N=11, 10, 11 & 10 mice, AF, AM, MF and MM groups, respectively. Normality of each dataset was assessed using the Shapiro-Wilk test, Means shown with overall distribution, Error bars are +/-standard error of the mean.

## Discussion

Overall, our work revealed an interesting sexual dimorphism in innate visually triggered approach versus arrest orienting that arises during adolescence in mice that accounts for significant differences in orienting reaction times in the natural prey capture context. Sensitivity to prey and visual motion is elevated in adolescents relative to adults, yet female adolescents are more likely to immediately approach and/or rapidly shift from arrest responses to approach relative to adolescent males, which are more likely to immediately arrest leading to slightly elevated approach times towards live crickets relative to their female peers (**Fig. 1 & 3, Supplemental Videos 1-8**). At the same time, there was marked reduction in predatory aggression in adolescent mice (**Fig. 2**). This suggests that in adolescence the circuits that link sweeping visual motion to either approach or arrest responses in mice may be flexibly controlled by neuromodulator systems^24^. Our work also provides additional evidence that “prey-like” visual motion is inherently salient even during development and drives rapid orienting behavior, consistent with previous studies demonstrating “sensory-triggered” predatory hunting controlled by specific subcortical circuits in mice^10^. However, in adolescents the link between visual motion detection and aggressive predatory responses is immature and appears differentially gated in the sexes by hunger state. Our data showing that adolescent males are more responsive than females to coupling their approach to attacks after just one night of food restriction supports that the two sexes differ in their baseline motivational state(s) that can be coupled to “sensory-triggered” predatory hunting perse. It is interesting to note that exposure to predatory threat in mice leads to sexually dimorphic effects on aggressive behavior and appetite^25^. As we scored attack attempts during prey exposure in addition to capture “success”/eating across days, we think it is most probable that differences in capture success mostly reflect significant differences in internal states related to arousal and/or hunger motivation rather than direct differences in the capability for perceptual or motor learning. Consistent with this idea, male C57BL6 mice undergo a significantly steeper gain in body weight and size relative to females starting at Postnatal day 35/5 weeks of age^26^. Thus, early adolescence may be a unique time in the life span of mice where the two sexes are differentially tuning visuo-motor orienting behaviors to subserve distinct adaptive behavioral outcomes^21,24^.

Notably, sex differences in diverse home cage behaviors such as nest building and saccharin preference are known to emerge during the onset of puberty in mice^21,24,27^. Some sexually-dimorphic behaviors with pubertal onset dissipate in adulthood, while others persist^21,24^. Our data demonstrate that sex-differences in innate orienting responses evoked by visual motion are specific to adolescence and indicate that each sex may go through a distinct developmental process that leads to more homogenous behavioral responses in the adult. In females, immediately approaching object motion would have to be more strongly inhibited, while arrest responses would have to be inhibited in males. This general idea would be consistent with a previous study showing that in male adult mice, hesitance and avoidance behaviors must be inhibited before attack and approach behaviors are optimally released in the presence of cockroaches^13^. Our own previous work, while it included both sexes, was biased towards a larger male population^14^. Interestingly, here, we find that adults of both sexes engage initially in alternating bouts of approach and “escape” when first encountering live crickets (**Supplemental Videos 1 & 2**). This “back and forth” response is significantly attenuated once food restriction begins and success capturing and eating insects occurs^14^. This might suggest then that females acquire their hesitance/inhibitory control over approach later in life. Further, the enhanced approach response in female adolescents that we observed is intriguing, but it remains unclear how females might specifically benefit from this bias during adolescence in nature. It is possible that it may give females an advantage in learning whether novel stimuli are threatening or rewarding early in life so that they can later quickly couple with predatory or defensive attack. A field study of birds adapting to urban environments found significant sex differences in “boldness”/ exploratory behaviors and avoidance behaviors that correlates with cortisol levels^23^. Further, one recent study shows that male and female adult mice have a different circuit configuration for responding to potential threats versus rewards routed from the ventral hippocampus to the Nucleus Accumbens^28^.

Female mice route both potential rewards and risks through the same circuit elements^28^. It is fascinating to speculate that the approach bias that we observe may help females refine and tune such circuitry during encounters with ambiguous novel stimuli that raise arousal levels. In this case, we speculate that they could gather additional sensory information that can optimally balance rapid responses to potential threats versus highly rewarding stimuli in the environment. Indeed, sweeping visual motion stimuli may be inherently ambiguous alone, despite a bias that they may represent potentially nutritious food^15^. However, there is so far no field, scene statistics nor ecology data to support speculation about how this wiring or another might be used to flexibly route natural stimuli to adaptive outcomes in a sex-specific way.

In future work, it will be interesting to determine which circuit elements may allow mice to flexibly link sweeping motion stimulus perception to approach, arrest or escape. We and others have already shown that adult mice of both sexes modulate between approach and escape when first encountering insects in a novel environment^13,16^ and that prey hunting experience increases approach probability of both sexes when they are adults^14^. In the context of threat responses, it is also known that the same visual input, an overhead looming stimulus, can evoke either an arrest or escape to shelter response mediated by distinct output channels^17^. In this case, the same visual information could either be routed to dorsal lateral geniculate nucleus (visual thalamus) to mediate rapid escape, or could be routed to parabigeminal nucleus to enhance arrest/freezing behavior^29^. We speculate that similar downstream targets may flexibly mediate differing rapid orienting responses in the predatory context, and, that these connections may be differentially recruited or subject to distinct neuromodulation in male versus female adolescents.

Another formal possibility for the enhanced approach responses and contact seen in adolescent mice overall could relate to a general, non sex specific, reduction in perceived risk in the prey capture and sweeping motion stimulus context in both sexes of adolescents^30^. Survival is not only dependent on the avoidance of threat, but of capitalizing on resources. Adolescence is characterized by an increase in risk-taking behavior^30^. Mice do not disperse from their mothers until late adolescence or early sexual maturity, thus practicing hunting skills during the early to mid-adolescence stage provides them with training for adulthood without the risk of starving if their efforts are unsuccessful. It is still unclear whether the live crickets and/or the virtual stimulus were perceived as possible threats. Though the adolescent mice did not exhibit thigmotaxis nor adopt postures that typically indicate anxiety in adults, the known reduction in threat response to occur in adolescence presents a confound for interpretation of this state via behavioral analysis alone^21^. Indeed, enhanced sensation seeking, exploration and impulsive responses to sensory stimuli in ambiguous or novel situations are thought to underlie human adolescent “risk-taking” behavior and are thought to be dissociable from “risk taking” behavior that emerges as a result of lacking higher-order risk assessment decision making under “known” risk conditions^30^. It is thought that one benefit that would outweigh the possible costs associated with enhanced risk-taking, regardless of its cause, at this stage of development is the ability to learn from the experiences themselves. Indeed, adolescence in mice is a period of heightened goal-directed learning and flexibility^31,32^. It would be important in future work to understand whether female adolescent mice benefit in other learning paradigms from their enhanced approach type visual responsiveness to external, salient stimuli^33^.

## Methods

### Resource Availability

Further information and requests for resources and reagents should be directed to and will be fulfilled by the Lead Contact, Jennifer L. Hoy.

### Materials availability

This study did not generate new unique reagents. However, a parts list and tips to create and run the described behavioral assays as well as complete datasets as raw movies and pose estimation model data can be obtained by request from the lead contact.

### Experimental model and subject details

All animals were used in accordance with protocols approved by the University of Nevada, Reno, Institutional Animal Care and Use Committee, in compliance with the National Institutes of Health Guide for the Care and Use of Laboratory Animals. Both male and female mice were used in this study. A total of 47 P30-45 adolescent mice and 53 P60-79 mature mice were used. The number of male and female subjects studied are similar. The specific number used in each statistical comparison is noted in figure legends. Mice were group housed with up to 5 animals per cage in an on-campus vivarium with *ad libitum* access to water and food (Envigo, Teklad diet, 2919), except where noted that mice experienced food restriction. The vivarium was maintained on a 12 hour light/dark schedule, and all testing occurred within 3 hours of the dark to light transition. Food restriction consisted of removing the food hopper from cages just before the light to dark transition (before the entry to dark phase) and testing animals just after the dark to light transition, 12-16 hours after food was removed from the homecage.

After hunting, mice were allowed to return to home cage with ad libitum access to their food hopper during the rest of their light phase cycle until it was once again removed prior to entry into the dark phase of their light cycle. Body weight measurements were taken to ensure that no participants lost more than %15 of their initial baseline weights.

## Method Details

### Apparatus

The cages of mice used for experimentation were transferred from the vivarium into a light-proof, double-wall sound room in order to control environmental conditions. During all data acquisition sessions, the door to the room was closed and the lights were turned off. For each session, each mouse was individually removed from its home cage and placed into a square, white acrylic open field arena with white vinyl flooring, 61 cm long x 61 cm wide x 3 cm high. One set of opposing sides are white acrylic walls and the other two opposing sides are composed of Hewlett Packard VH240a video monitors measuring 60.5 cm diagonally, with a vertical refresh rate of 60 Hz and 1920×1080 pixel resolution. A solid white background was projected on each monitor in order to display even lighting throughout the arena which measured 250 cd/m_2_ of luminance. One of the monitors was used to display the visual stimulus, which consisted of a small black ellipse, 2 × 1 cm, generated by a customized MATLAB Psychophysics toolbox script, which was programmed to move back and forth horizontally across the monitor with the center of the stimulus consistently 2.5 cm above the floor. A Logitech HD Pro Webcam C920 digital camera was placed suspended overhead to record mouse and stimulus positions at 30 frames per second throughout each session.

### Visual Stimuli

Visual stimuli were generated with MATLAB Psychophysics toolbox (Brainard, 1997) ^34^and displayed on an LCD monitor (60 Hz refresh rate, ∼50 cd/m2 luminance) in a dark room. The computer monitors replaced two sides of a 4-sided behavioral arena. To mimic insect proportions, we displayed ellipses with a major horizontal axis that was 2 times the size of the minor vertical axis and displayed stimuli that were 2cm along the horizontal axis as these stimuli evoked the most frequent approaches in adult mice_9_. For a presented stimulus with a major axis of 2 cm, this corresponds to the presentation of a 4°sized target from 20 cm away from the screen. Stimulus speed was kept constant at 2 cm/sec as this speed evoked the most approach and least amount of arrests in adult mice as previously reported in Proacci et al., 2020^9^. This meant that a single stimulus sweep lasted for approximately 30 sec on the screen.

### Behavior Baseline Mice

Prior to testing, mice were acclimated to handlers and the behavior arena for 2 days: days 1 and 2 included three 3-minute handling sessions; on day 2, mice were exposed to the behavior arena for 3 5-minute sessions each. The arena floor was cleaned thoroughly with 70% EtOH after each mouse was removed to mitigate odor distractions. Testing did not begin until the present mouse moved throughout the arena, away from walls, and demonstrated self-grooming behaviors, which indicated behaviors least associated with anxiety and fear. Mice in the cohort who did not meet these criteria by the end of the third day prior to testing received an extra day of handler and arena acclimation identical to day 2.

On testing day, each mouse was placed in the arena for a 1-minute habituation period. This period was used as a control trial to assess mouse behavior in the absence of a stimulus. After this period, the stimulus was presented on the display monitor described above. The virtual stimulus moving 2 cm/s was presented for 60 seconds.

For live-prey exposure, each mouse was placed into the arena with a live cricket for a total of 3 rounds, up to 5 minutes each. If the mouse caught the cricket within this period of time, it was given up to a total of 3 crickets during that 5-minute time window. If it did not catch the cricket within 5 minutes, the mouse was returned to its home cage and the arena was cleaned in preparation for the next mouse to be tested. After testing, all mice were returned to their home cages with standard food. Each session was recorded as a separate video for a total of 5 videos per mouse on test day (1m habituation, 1m virtual stimuli, & 5m live prey capture).

### Data Analysis

DeepLabCut was used to digitize and extract 2-dimensional coordinates of the mouse’s nose, two ears and body center, as well as the center point of the stimulus (cricket or ellipse stimulus), throughout the video recordings at 30 frames per second. These tracks were entered into customized MATLAB scripts to extract behavioral parameters such as mouse and stimulus/cricket speed, stimulus/cricket angle, and range between mouse and stimulus/cricket.

An arrest was defined as any time the mouse’s nose and body moved less than 0.5 cm/sec for a duration of 0.5 - 2 seconds. Arrests that occurred in the absence of a visual stimulus, or when the stimulus was more than 130 degrees from the bearing of the nose, were excluded from analysis of visually-driven arrest responses. An approach was defined as any time the mouse’s nose came within 4 cm of the stimulus center after moving toward the stimulus for a distance of at least 5 cm, and at an average approach speed of at least 15 cm/sec. Using these definitions, we computed the percentage of stimulus trials in which each behavior was observed, as well as the number of arrests and approaches that occurred during individual trials.

Statistics were performed using MATLAB software. Where means are reported, ANOVA or repeated measures ANOVA were used to identify main effects followed by posthoc testing with correction for multiple comparisons when identifying the specific significant differences between groups. Where medians are reported, the non-parametric Friedman’s test was used followed by rank sum. Where percentages are compared, Fisher’s exact test was used. Test results with a p-value of < 0.05 were considered significant.

## Acknowledgements

We would like to thank undergraduate researchers Suvrajyoti Rout, Stetson Necaise and Aryanna Ortega for their support in handling experimental mice and manually confirming behavioral responses to visual stimuli in mice. This work funded by RO1 EY032101-01A1 to PI Hoy.

## Notes

### Competing Interest Statement

The authors have declared no competing interest.

